# *erm-1* mRNA and ERM-1 protein co-translationally localize to the plasma membrane through a microtubule- and BMK-1-dependent pathway

**DOI:** 10.64898/2026.05.15.725403

**Authors:** Naly Torres Mangual, Karissa Coleman, Erin Osborne Nishimura

## Abstract

The Ezrin, Radixin, and Moesin (ERM) family of proteins anchors the actin cytoskeleton to the plasma membrane for the purpose of either stabilizing or altering cell shape. In *Caenorhabditis elegans*, ERM-1, is essential for cell polarity, signaling, intestine development, and larval viability. Interestingly, ERM-1 proteins are produced by *erm-1* mRNA transcripts that concentrate at the plasma membrane in embryos. The localization of *erm-1* mRNA to the plasma membrane occurs in a 3’UTR-independent, translation-dependent manner, directed by the PH-subdomain within ERM-1’s N-terminal FERM domain. This has led to the model that *erm-1* mRNA, its associated ribosome, and its emerging nascent peptide are all transported together to the plasma membrane as a complex. Here, we characterize the transport mechanism. Using a microscopy approach, we observed that the localizations of *erm-1* mRNA and ERM-1 protein to the plasma membrane were disrupted by nocodazole treatment, illustrating a microtubule role. Furthermore, *erm-1* mRNA and ERM-1 protein localized to the plasma membrane independently of myosin and dynein motors, but dependent on the kinesin *bmk-1 (bmk-1)*, a plus-end-directed, Kinesin-5 family motor protein. Loss of *bmk-1* did not reduce the total number of *erm-1* mRNA molecules in the cell, arguing against a diffusion- and protection-based mechanism of mRNA localization. Together, these findings suggest that *erm-1* mRNA is localized via an active transport pathway mediated by a plus-end-directed kinesin adapter. Interestingly, loss of *bmk-1* led to diffuse localization of ERM-1 protein along the plasma membrane and reduced ERM-1 protein levels at the site of abscission, the midbody, and the midbody remnant. This suggests that ERM-1 local translation at the plasma membrane is critical for its protein’s ultimate spatial patterning in the cell.

## INTRODUCTION

Gene expression dynamics drive embryogenesis by orchestrating cellular differentiation and growth. Because *de novo* transcription is paused in early embryos, gene expression regulation is often achieved through post-transcriptional mechanisms acting on maternally inherited mRNAs (Edgar et al., 1994; Guven-Ozkan et al., 2008; Tadros and Lipshitz, 2009; Vastenhouw et al., 2019). Recent work in *Caenorhabditis elegans* has shown that many post-transcriptionally regulated maternal mRNAs concentrate at interesting locales within the cell, such as at membranes, condensates, or pericentriolar material (Parker et al., 2020; Winkenbach et al., 2022). This presents an opportunity to study the relationships between post-transcriptional regulation and mRNA localization, yielding insights that are often applicable beyond embryos.

The phase of development from oocyte to early embryo, termed the oocyte-to-embryo transition, has long been recognized for its high propensity for mRNA localization. In Xenopus oocytes, numerous mRNAs localize to the vegetal or animal hemispheres where they contribute to primary germ layer development, axis formation, and cell differentiation (Mowry and Cote, 1999). In *Drosophila* syncytial embryos, antipodal localizations of *bicoid* (anterior), *oskar* (posterior), *nanos* (posterior), and *gurken* (posterior and anterior-dorsal) mRNAs generate morphogen gradients that determine axis formation and polarity (Driever et al., 1989; Lasko, 2012; Lu et al., 2020; Nüsslein-Volhard et al., 1987). Indeed, up to 70 % of *Drosophila* embryonic mRNAs are estimated to be localized, but most have unclear consequences on development (Lécuyer et al., 2007; Lécuyer et al., 2009). In *C. elegans*, a cohort of mRNAs concentrates within germline-associated condensates (P granules and germline P-bodies), structures that are preferentially inherited and stabilized in the progenitor germ lineage (Cassani and Seydoux, 2024; Lee et al., 2020). Of these, *nos-2* mRNA is required to localize to P granules for germ cell health and survival, but other P granule-localized transcripts, whose patterning results indirectly from translational repression or decay, are not required for germline development (Lee et al., 2020; Putnam et al., 2023; Scholl et al., 2024). One common feature among the examples above is their dependence on cis-regulatory sequences housed in either the 3’ Untranslated Regions (3’UTRs) or arising from key splicing events. These signals are necessary (and often sufficient) to direct mRNA localization (Jadhav et al., 2008; Mayr, 2019; Mukherjee et al., 2019; Oldenbroek et al., 2012). Though the detailed mechanisms of transport can vary, most of these examples converge on a core model in which sequences within the mRNA (also known as zipcodes) are recognized by effector proteins, then linked to cytoskeletal motors, and then delivered to their destinations for ultimate translation or other fates (Das et al., 2021; Johnston, 2005).

In reality, the core model of 3’UTR-directed mRNA localization is only the tip of the iceberg. In recent years, numerous examples of mRNA localization across a wide range of organisms have yielded deeper insight into the varied ways translation, mRNA decay, RNA interference, condensate biology, and development all connect to mRNA localization (Chouaib et al., 2020; Keiper and Huggins, 2025; Liao et al., 2019; Ma and Mayr, 2018; Müntjes et al., 2021; Parker et al., 2022; Safieddine et al., 2025). Among these novel mechanisms, is a new paradigm in which the nascent peptide directs the transport of its associated ribosome and mRNA, moving the ribonucleoprotein complex (RNC) to specific subcellular destinations (Chouaib et al., 2020; Tocchini et al., 2021; Winkenbach et al., 2022). This 3’UTR-independent, co-translational mRNA localization mechanism is conceptually analogous to, but mechanistically distinct from, the SRP pathway that targets secreted or transmembrane proteins to the rough endoplasmic reticulum (Walter and Blobel, 1981). To date, co-translational transport has been observed at multiple cellular locations, directing mRNAs to the plasma membrane (Chouaib et al., 2020; Tocchini et al., 2021; Winkenbach et al., 2022), nuclear membrane (Chouaib et al., 2020; Lautier et al., 2021; Parker et al., 2020), mitochondria (Chang et al., 2025; Fazal et al., 2019; Müntjes et al., 2021; Tian et al., 2019; Zhu et al., 2025), and centrioles (Ryder and Lerit, 2018; Sepulveda et al., 2018; Zein-Sabatto et al., 2024).

Despite evidence that co-translational mRNA localization is widespread, its functional impact on protein, cellular, and developmental outcomes remains unclear. An emerging view is that local translation of key membrane-associated proteins may be beneficial for producing pools of proteins in response to local signals and without the need for further transport (Du et al., 2007; Lécuyer et al., 2009). Another idea is that local translation may facilitate protein-protein interactions and large complex assembly (Mayr 2018; Engel et al. 2020; Horste et al. 2023).

One example of co-translational mRNA localization in *C. elegans* embryos is *erm-1* (Parker et al., 2020; Winkenbach et al 2022). ERM-1 belongs to the conserved Ezrin/Radixin/Moesin family of proteins that physically link the plasma membrane to the actin cytoskeleton. In *C. elegans*, ERM-1 is the sole homolog of this family and plays essential roles in cell polarity and morphogenesis. Loss of ERM-1 function leads to severe defects in epithelial organization, particularly in intestinal development, resulting in larval lethality (Furden et al., 2004; Göbel et al., 2004; Ramalho et al., 2020). In 2022, Winkenbach et al. reported that *erm-1* mRNA localization at the plasma membrane was translation-dependent and required the protein’s N-terminal FERM domain for targeting (Winkenbach et al., 2022). The FERM domain binds PIP_2_ phospholipids within the plasma membrane through the coordinated efforts of four lysine residues in its Plekstrin Homology (PH) subdomain. In contrast, ERM-1’s C-terminal Ezrin-Radixin-Moexin Association Domain (C-ERMAD) binds the actinomycin network (Fehon et al., 2010). ERM-1, therefore, bridges the plasma membrane and the internal actinomyosin cytoskeletal structure. Work in other organisms suggests that ERM family proteins can influence cell shape by stiffening the cortex or by promoting protrusion formation to generate microvilli or blebs.

It is unclear why the cell translates ERM-1 proteins locally at the membrane. However, Van der Salm et al., recently demonstrated that local translation is required for full ERM-1 function. To illustrate this, they tethered *erm-1* mRNA to the nuclear membrane, verified that ERM-1 proteins could still be translated at their new location, and reported disrupted intestinal development (Salm et al., 2025a). This illustrates that local translation of ERM-1 is essential for its function and is purposefully regulated.

Here, we investigate the mechanisms governing the localization of *erm-1* mRNA and ERM-1 protein to the plasma membrane in *C. elegans* early embryos and during mid-stage intestinal development. Using fixed imaging approaches, we assessed the cytoskeletal components and motors required for *erm-1* mRNA localization. We provide evidence that the localization of the erm-1/ERM-1 translating complex depends on microtubules and a specific kinesin, *bmk-1*. Altogether, our work aims to define how co-translational targeting is coordinated to ensure proper embryonic development.

## RESULTS

### *erm-1* mRNA localization is microtubule-dependent

A variety of mechanisms have been characterized that orchestrate sequence-directed mRNA transport. These mechanisms broadly parse into three classes: active transport, diffusion (often coupled with anchoring), and mRNA decay (Das et al., 2021; Lashkevich and Dmitriev, 2021; Martin and Ephrussi, 2009). The three classes are not mutually exclusive; in fact, multiple mechanisms can work together on a single target mRNA. For cases in which mRNA localization occurs co-translationally through peptide-encoded signals, fewer known mechanisms have been described. One is the SRP-directed pathway that directs the co-translational transport of secreted and transmembrane proteins to the endoplasmic reticulum, where they can either be inserted into the membrane or packaged into vesicles (Ast et al., 2013; Chartron et al., 2016). In this pathway, the SRP particle (a protein-RNA complex) binds an emergent N-terminal “signal sequence”, stimulates translational elongation arrest, and then diffuses to the endoplasmic reticulum, where it docks with an SRP receptor. In other cases, translating mRNAs can hitchhike on organelles such as lysosomes, endosomes, or mitochondria to reach their destinations (Liao et al., 2019; Müntjes et al., 2021; Nagano and Araki, 2021).

The mode of ERM-1 co-translational transport to the membrane is unknown. Live images of *erm-1* mRNA using a PP7 reporter system showed both non-directed and rapid, directed movements, supporting a hypothesis that ribosome-free *erm-1* mRNA moves in non-directed movements, whereas ribosome-bound *erm-1* mRNA moves via active transport (Li et al., 2021). However, live images of actively translating ERM-1, using the SUNTAG system, also observed both types of movement, illustrating that ribosome-bound ERM-1 can move in both non-directed and directed fashions (Salm et al., 2025a; Salm et al., 2025b). Therefore, the mechanism of ERM-1 transport is an open question, but the observance of rapid, directed motion to the plasma membrane in both PP7 and SUNTAG systems, suggests that active transport, mediated through either microtubules or actin, has a role.

To test whether *C. elegans erm-1* localization to the plasma membrane depends on microtubules, we treated TBB-2::GFP (tubulin GFP) worms with Nocodazole and assessed the resulting embryos for *erm-1* localization by smFISH. Because small-molecule drugs do not readily diffuse into *C. elegans* embryos due to their permeability layer, we first depleted that layer through PERM-1 (permeability barrier defective) depletion using RNAi (Olson et al., 2012). This approach typically achieves a population of >75% permeabilized embryos with minimal deleterious effects (Carvalho et al., 2011).

To disrupt microtubules, we treated the permeabilized TBB-2::GFP embryos with 0 μM Nocodazole (DMSO control) or 150 μM Nocodazole for 15 minutes. Nocodazole is a potent microtubule-depolymerizing drug. The TBB-2::GFP reporter confirmed that microtubules remained intact in control embryos but were effectively disrupted following nocodazole (Fig. 1B). Nocodazole treatment resulted in a dramatic depletion of *erm-1* mRNA localization at the cell-cell junctions of 2-cell stage embryos, as evidenced by smFISH (Fig. 1C, D, E). In contrast, the homogenously distributed *set-*3 mRNA remained unchanged. Notably, the *erm-1* mRNA was not uniformly distributed between AB and P_1_ cells upon treatment; instead, it remained enriched in AB cells, suggesting that cell-specific patterns of *erm-1* mRNA enrichment do not require its localization (Tintori et al., 2016). Strikingly, we observed no statistically significant differences in total mRNA abundance between DMSO and Nocodazole-treated animals (Fig 1F). These data suggest that *erm-1* mRNA is not dependent on localization to guard against degradation, as mRNA abundance did not decline when localization was disrupted.

**Fig 1.**
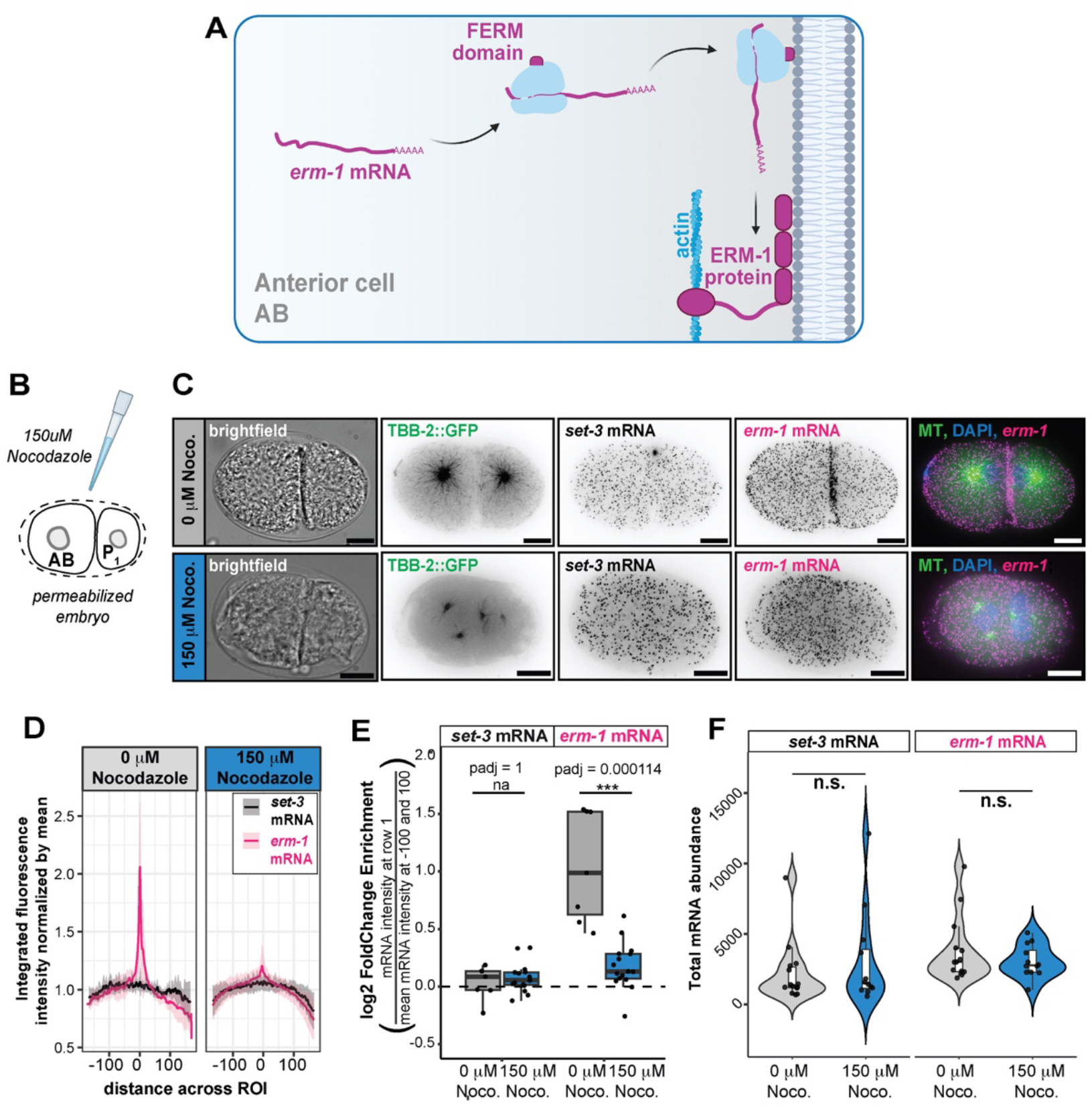
Microtubules are required for *erm-1* mRNA localization to the plasma membrane. **(A)** A model of *erm-1*/ERM-1 co-translational localization to the plasma membrane based on evidence that *erm-1* and ERM-1 movement to the membrane requires active translation and ERM-1’s own N-terminal FERM domain. **(B)** Embryos were permeabilized by growth under *perm-1* RNAi depletion then soaked in Nocodazole to depolymerize microtubules. **(C)** Representative 2-cell embryos depicting single-slice brightfield illumination, maximum intensity projections of TBB-2::GFP, a microtubule marker (green), maximum intensity projections of smFISH for *set-3* mRNA symmetrical control (black) and *erm-1* mRNA (magenta), and merged images.Embryos were treated with 0 μM Nocodazole DMSO control (top) or 150 μM Nocodazole for 10 minutes (bottom). Scale bars 10 μm. **(D)** Integrated fluorescence intensity of *set-3* and *erm-1* mRNA molecules across a rectangular region spanning the anterior-posterior body axis on sum-projected images. **(E)** Values from **D** were analyzed by taking the log_2_ mRNA signal intensity ratios at “distance across the ROI” = 1 divided by the mean of “distance across the ROI” = -100 and 100 to illustrate the peak versus background fold change enrichment. Statistics were calculated using Wilcoxon ranked sum tests and BH-adjusted p-values (p-adj) are shown. **(F)** Violin plots showing differences in the total abundances of *set-3* mRNA (Wilcox statistical test p-value = 0.6251) and *erm-1* mRNA (Wilcox statistical test p-value = 0.4494) mRNA between DMSO control and Nocodazole treated embryos. control n = 15, 150 mM nocodazole n = 18, 2 reps each.

### Dynein is required for full *erm-1* mRNA abundance but not its localization

Microtubule-associated cargo transport can be mediated through two different classes of motor complexes, dyneins and kinesins. Dyneins traffic cargo largely in a minus-end direction, moving cargos to the center of the embryo and towards the centrioles (Canty et al., 2021). In contrast, kinesins typically transport their cargos to the plus ends of microtubules, trafficking cargo from the center to the periphery of the cell (Siddiqui, 2002; Yildiz, 2025). Though kinesins largely travel in the direction of *erm-1* mRNA transport, previous studies found that *erm-1* mRNA localization in adult intestines is dynein-dependent. That is, *erm-1* mRNAs coalesced into punctate droplets upon *dhc-1* RNAi in polarized, adult intestinal cells, as observed using the PP7 mRNA live imaging system (Li et al., 2021). To determine whether early embryos also utilize dynein for *erm-1* mRNA localization, we depleted dynein using *dhc-1* RNAi in a membrane marker strain (*ph::gfp)* and observe *erm-1* mRNA localization by smFISH.

The cytoplasmic dynein heavy chain 1, *dhc-1*, encodes a microtubule-associated motor protein required for retrograde transport, mitotic spindle assembly, nuclear positioning, and embryonic development in *C*.*elegans* (Schmidt et al., 2005). Consistent with these roles, we observed that *dhc-1*-depleted embryos exhibited defects in nuclear positioning, mitosis (Fig. 2A,B), and embryos did not progress properly through development. For these reasons, the inclusion of a membrane marker (PH::GFP) was essential to mark the location of membranes as DAPI staining and brightfield imaging were insufficient to detect them. To test whether the enrichment of *erm-1* or *set-3* mRNA at membranes had changed upon DHC-1 depletion, we calculated the fold-change between the density of mRNA at the membrane versus in the embryo as a whole (Fig. 2C). We found that of *erm-1* mRNA enrichment at membranes remained unchanged after dynein depletion despite the gross morphological changes to the membrane (Fig. 3B). However, the overall abundance of both transcripts was reduced upon dynein depletion (*set-3* mRNA, p-value = 2.904 e-06 and *erm-1* mRNA, p-value *=* 1.555036e-09) (Fig 3D). This suggests that although dynein was not required for *erm-1* mRNA localization to membranes in embryos, it was required for wild-type mRNA abundance.

**Fig 2.**
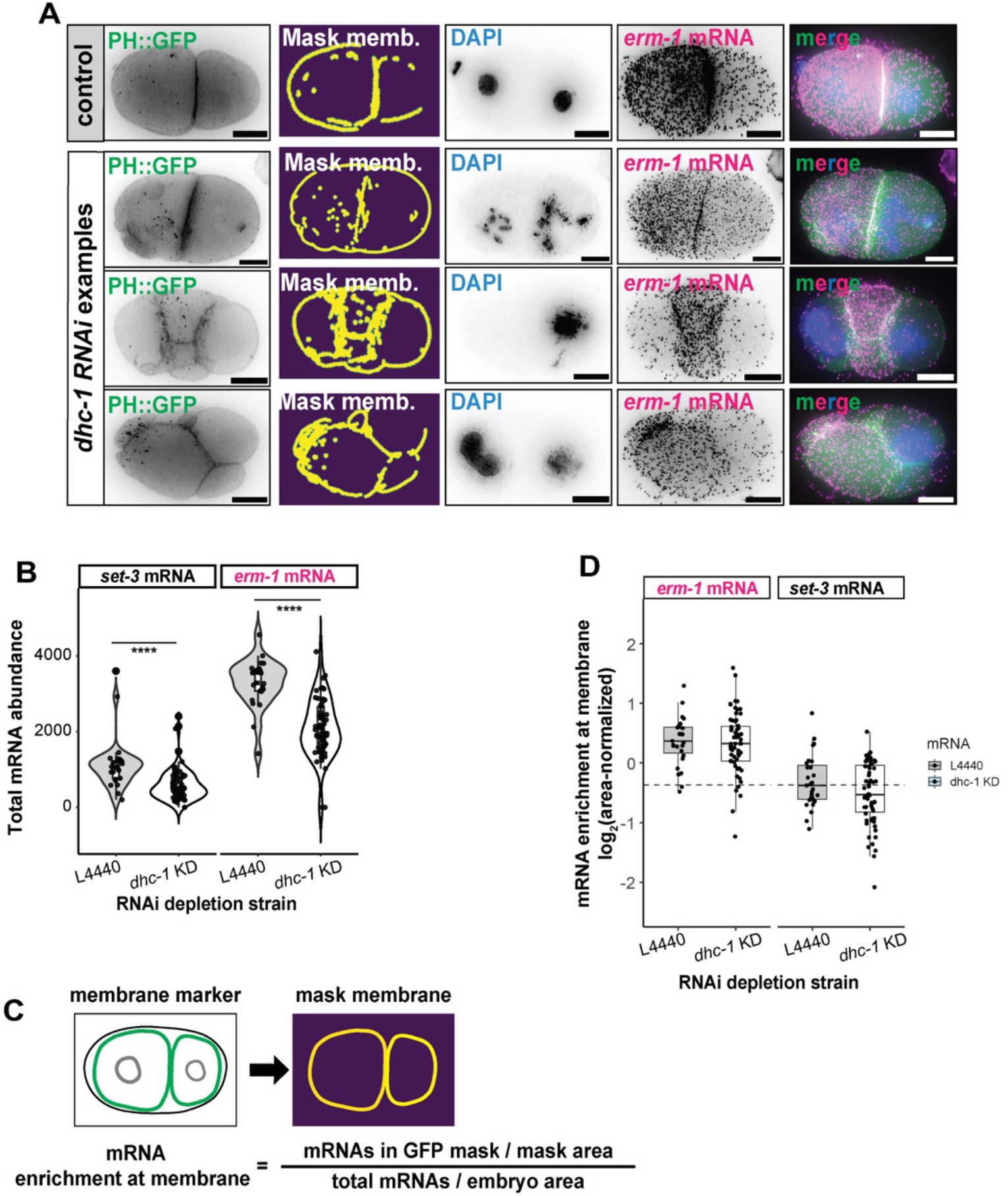
Dynein depletion in embryos affects overall mRNA abundance but not *erm-1* mRNA localization. **(A)** Representative micrographs of 2-cell stage embryos showing membranes (*ph::gfp*), DAPI showing abnormal nuclear divisions and *erm-1* mRNA smFISH under L4440 control or DHC-1 RNAi conditions. Merge of *erm-1* (magenta), membrane (green) and DAPI (blue). A range of phenotypes resulted from DHC-1 depletion. Three examples are shown of increasing severity. Scale bars 10 mM (**B**) Violin and box plot with jittered data points showing total abundance of *set-3* mRNA and *erm-1* mRNA in control L4440 or DHC-1 RNAi depletion (Tukey multiple comparisons of means *set-3* p-value = 2.904 e-06 ; *erm-1* p-value = 1.555036e-09). N-values ranged from 23-60 per condition across 3-4 reps. (**C**) mRNA density at membranes for each embryo was calculated by dividing the number of mRNAs in gfp mask normalized by mask area, divided by the total amount of mRNAs in the entire embryo.

**Fig 3.**
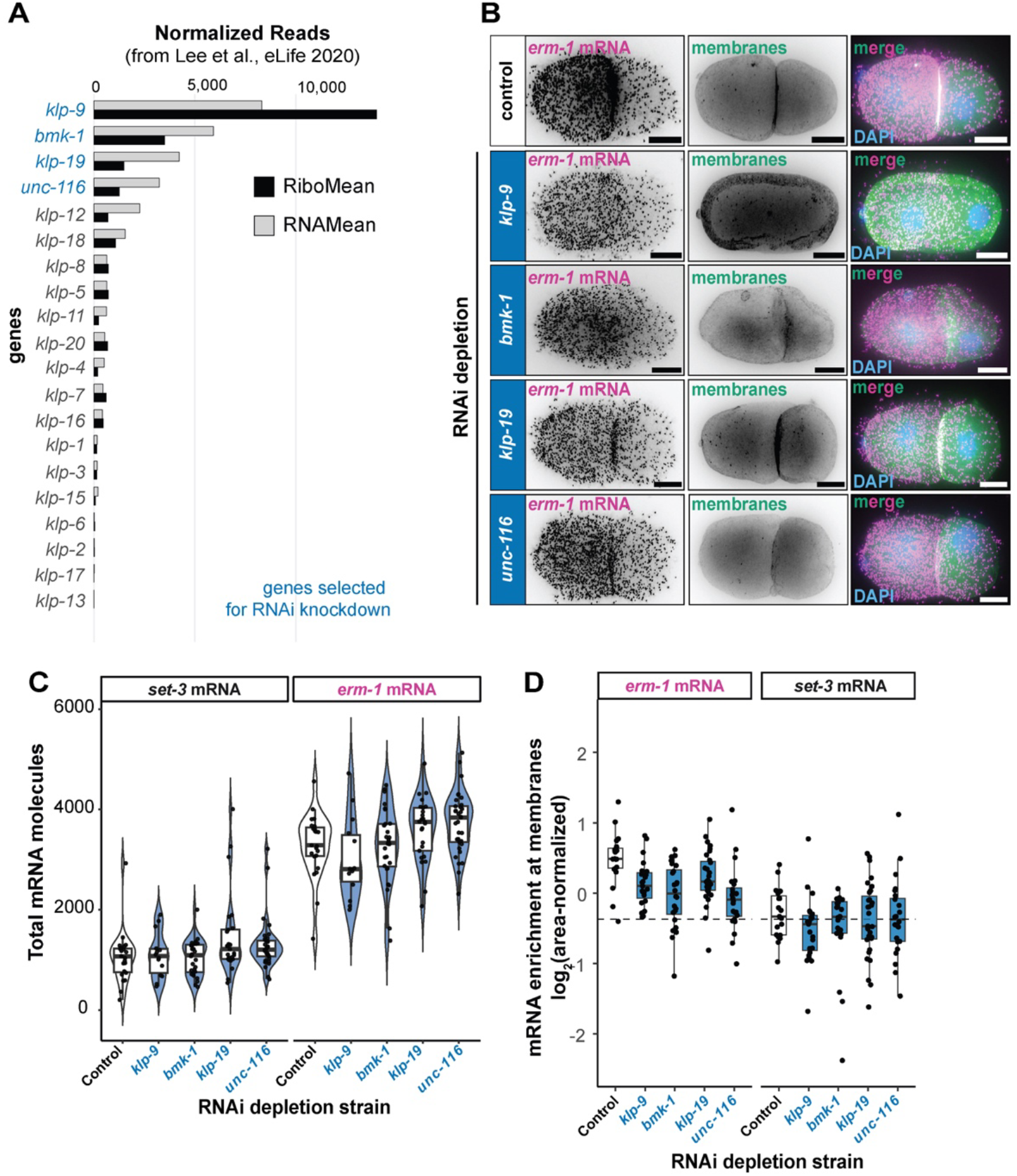
The kinesin BMK-1 is required for *erm-1* mRNA localization to the membrane of early embryos. **(A)** RNA-seq and ribosome profiling (black) and RNA-seq (gray) mean read counts generated from mixed-stage early *C. elegans* embryos (Lee et al., eLife 2020; GEO accession: GSM4148080). Four genes (blue text) were selected for a reverse genetic screen to assess their impact on *erm-1* mRNA localization. (**B**) Micrographs of early *C. elegans* embryos depleted for kinesin and assayed using smFISH for *erm-1* mRNA localization.Representative images of 2-cell stage embryos. *erm-1* mRNA(magenta), membranes (PH::GFP, green) and nuclei (DAPI, blue) are shown as individual color channels and merged. Scale bars 10 μM. (**C**) Quantification of total mRNA molecules from the full dataset depicted in **B** graphed as box plots, violin plots, and jittered data points. *erm-1* (right panel) and *set-3* mRNA (left panel), a homogenously distributed cytoplasmic control, are both shown. Empty vector RNAI control (L4440 plasmid) or kinesin depletion conditions are plotted. N-values range between 24-41 per condition across 3-4 reps. Significance was calculated using the Wilcox statistical test. (**D**) The enrichment of *erm-1* mRNA at membranes versus within the total embryo as calculated in Fig 2C,D.

The observed reduction in mRNA abundance could be the indirect result of broader defects in embryonic development. Indeed, the embryos we categorized as 2-cell- or 4-cell-stage embryos occasionally had supernumerary nuclei or were anucleated, suggesting pleiotropic, diverse phenotypes. Consistent with this, cells that had supernumerary nuclei had lower levels of mRNA, whereas anucleated cells accumulated a greater abundance of maternal transcripts. This observation suggests that dynein-dependent processes indirectly influence mRNA levels, potentially through pleiotropic effects including mitotic nondisjunction.

### BMK-1 is required for *erm-1* mRNA localization to the plasma membranes of 2-cell stage embryos

To identify microtubule-associated motors that may mediate *erm-1* mRNA transport, we next examined the kinesin family. While a small group of kinesins moves products toward the minus end of microtubules near centrosomes, most kinesins transport anterograde, towards the rapidly growing positive end of microtubules near the plasma membrane (Yildiz, 2025). The *C. elegans* genome encodes 21 kinesins (Siddiqui, 2002). To identify kinesin motors that would be likely candidates for *erm-1* mRNA transport, we assessed the expression and translation levels of each using previously published ribosome profiling and RNA-seq datasets from mixed-stage early embryos (Lee et al., 2020) (Fig. 3A). We selected the four most highly expressed kinesins: *klp-9, bmk-1, klp-19* and *unc-116* to use in a reverse genetics approach to survey for impact on *erm-1* mRNA localization (Fig. 3A, depicted in red).

Of the four kinesins depleted (*klp-9, bmk-1, klp-19* and *unc-116*), both *klp-9* and *bmk-1* yielded abnormal *erm-1* mRNA localization in 2-cell stage embryos (Fig. 3B, D). These embryos lost the classical *erm-1* mRNA patterning in which mRNA molecules are most abundant at the interface between the anterior AB and posterior P_1_ cells (Fig. 3B). It should be noted that *erm-1* mRNA shows the greatest enrichment at this face. They are still enriched at other membrane faces, but the curved surfaces of those membranes yield less dramatic enrichment after max projection. Though both *klp-9* and *bmk-1* yielded abnormal *erm-1* distribution, the impact of *klp-9* appeared indirect as membranes were missing at the AB/P1 interface. In contrast, RNAi depletion of *bmk-1* resulted in a specific loss of *erm-1* mRNA enrichment at the plasma membrane despite largely intact membrane morphology (Fig 3B, D). Notably, total RNA content of *erm-1* and *set-3* control mRNA was not altered when individual kinesins were depleted (Fig 3C). Depletion of *klp-19* or *unc-116* did not affect overall embryo morphology, and these embryos appeared wild-type on the surface until hatching. Altogether, these data suggest that bmk-1 is required for the enrichment of *erm-1* mRNA at the plasma membrane.

### BMK-1 is required for proper ERM-1 protein localization in intestinal cells

How does local translation of *erm-1* at the plasma membrane impact its encoded protein? To test this, we assessed the impact of *bmk-1* depletion (and its resulting loss of *erm-1* mRNA localization) on ERM-1 protein accumulation in BOX213 (ERM-1::GFP) worm strains. In wild-type, 2-cell stage embryos, ERM-1 is seen at the interface between the AB and P_1_ cells with peak accumulation at the site of previous abscission and in the midbody remnant. These two locations represent the last points of contact between dividing daughter cells undergoing cytokinesis and are the termination points for plus-end microtubules. Therefore, they often accumulate plus-end-directed cargoes. We found that when the kinesin BMK-1 was depleted, ERM-1 protein at the membrane was reduced and diffuse (Fig. 4A). That is, ERM-1 proteins spread out along membranes at the AB and P_1_ interface and failed to achieve a peak of localization at the site of previous abscission or in the midbody.

**Fig 4.**
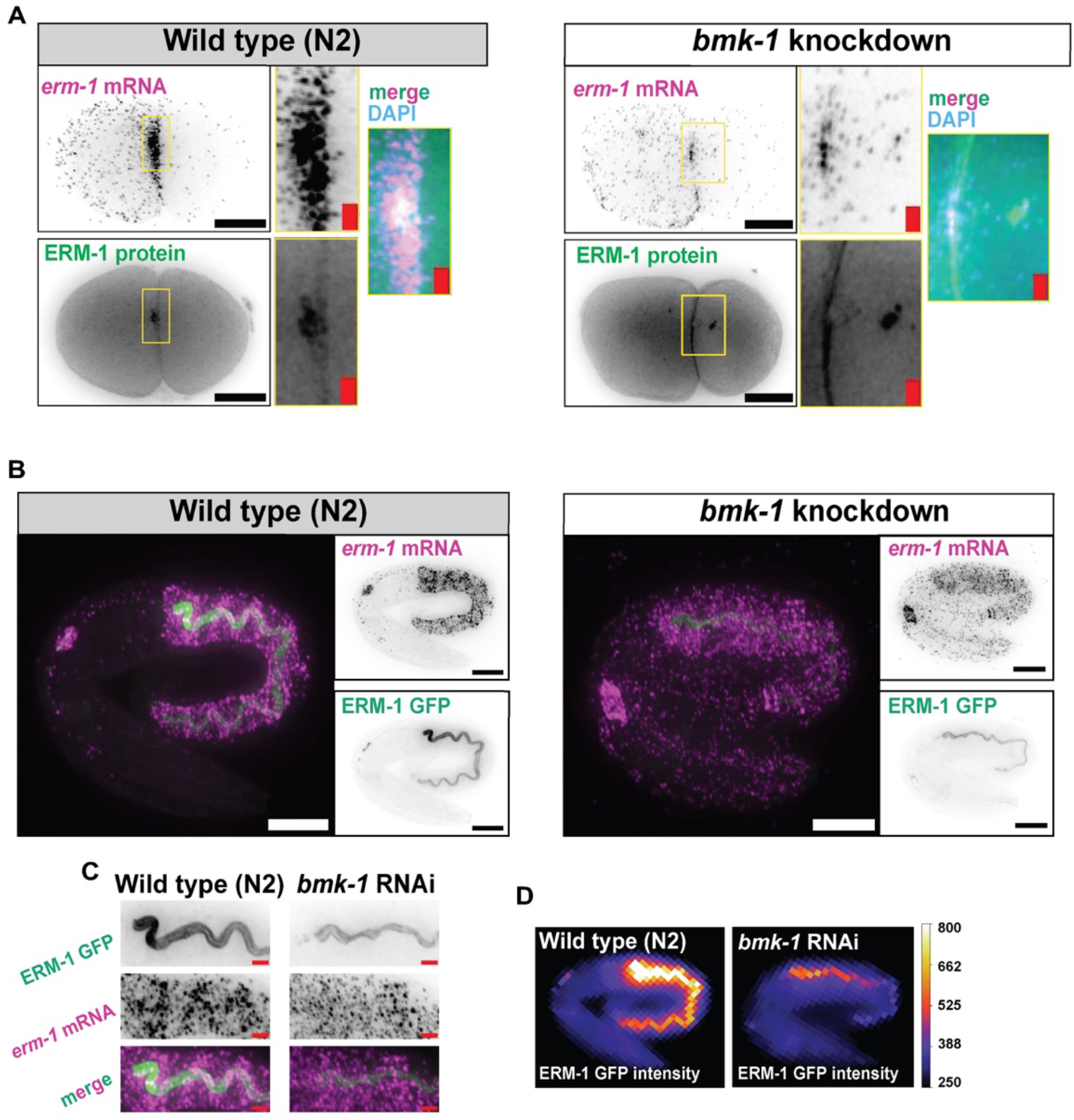
The *erm-1* mRNA and ERM-1 protein complex depend on *bmk-1* for directed localization. **(A)** Representative 2-cell stage embryos with smFISH probes for *erm-1* mRNA under L4440 (empty vector control) or kinesin *bmk-1* RNAi depletion conditions. *erm-1* mRNA (top) and ERM-1 protein from *erm-1::gfp* strain. Insets scale bar are 1 μm. (L4440 n = 44; *klp-14* n= 21, between 2-3 replicates. **(B)** Representative 2-fold embryo under L4440 (empty vector control) or kinesin *bmk-1* RNAi depletion conditions with smFISH probes for *erm-1* mRNA (top). ERM-1 protein (bottom) as GFP fluorescence from the *erm-1::gfp* strain. (control n=21; *klp-14* n= 42 between 2 replicates **(C)** Inset of anterior intestine from embryos shown in **B**. Intestine insets scale bar in red are 2μm. **(D)**. Fluorescence intensity heatmaps of intestines show in **B**. Heatmaps were generated with standardized scales and intensity thresholds to show direct comparison between ERM-1 abundance in each condition.

In *C. elegans*, the local translation of ERM-1 at the plasma membranes of intestinal cells is required for proper ERM-1 protein accumulation and proper intestinal development. Therefore, we next asked how BMK-1 depletion affected ERM-1 protein levels in intestinal epithelial cells. In wild-type intestines, ERM-1 accumulates at the apical lumen of embryonic, larval, and adult intestines and serves as a common marker defining this feature. Interestingly, previous work has shown that the *erm-1* mRNA localizes to all faces of the plasma membrane of intestinal epithelial cells (even the basolateral face where ERM-1 protein is low). This localization difference between the *erm-1* mRNA (basolateral and global) and the ERM-1 protein (apical) has given rise to the Two-Step Hypothesis, which posits that ERM-1 proteins undergo post-translational migration to the apical face.

BMK-1 depletion resulted in a marked reduction of ERM-1 protein at the apical lumen of the developing intestine (Fig 4B, C, D). *erm-1* mRNA accumulation at the plasma membrane was also diffused. The morphology of the intestine was altered, showing fewer turns and crenulations than the wild type. BMK-1-depleted worms survived until hatching. Together, these results support a model in which BMK-1-dependent *erm-1* local translation serves to direct both co-translational production of ERM-1 and its post-translational migration to the apical face of intestinal epithelial cells.

## DISCUSSION

Here, we illustrate that *erm-1* mRNA localizes to the plasma membranes of *C. elegans* embryonic cells through an active transport mechanism that utilizes microtubules and the plus-end-directed kinesin, BMK-1. These findings expand the model of ERM-1 co-translational transport by eliminating diffusion and mRNA decay as possible mechanisms and by identifying a key kinesin factor.

### A more detailed model of ERM-1 co-translational transport

By incorporating our findings into previous reports of *erm-1* behavior, a more detailed pathway emerges (Fig. 5). In this pathway, *erm-1* mRNAs initiate translation in the cytoplasm to produce an N-terminal FERM peptide (Salm et al., 2025a; Winkenbach et al., 2022). This peptide is likely recognized by an effector of unknown identity that, in turn, recruits the kinesin BMK-1. The association of BMK-1 likely triggers the transport of the *erm-1*/ERM-1 translational complex along microtubules. The transported complexes are comprised of the *erm-1* mRNA, the ribosome, the nascent ERM-1 FERM peptide, and any associated proteins. They move towards the plus ends of the microtubules which terminate at the cellular periphery.Though we do not know whether translational pausing or re-initiation occurs in this pathway, we hypothesize a model in which ERM-1 translation elongation and termination occur at the membrane. One idea that will be explored in more detail below is that local translation may be important for ERM-1 to form proper contacts with the plasma membrane and to become phosphorylated at key residues associated with its active, open state (Ramalho et al., 2020; Sepers et al., 2022; Zhang et al., 2020). Once ERM-1 proteins have been produced at the plasma membrane, they undergo a post-translational migration event, bringing them to the site of abscission or the midbody in early embryonic cells, or to the apical face of intestinal epithelial cells.

**Fig 5.**
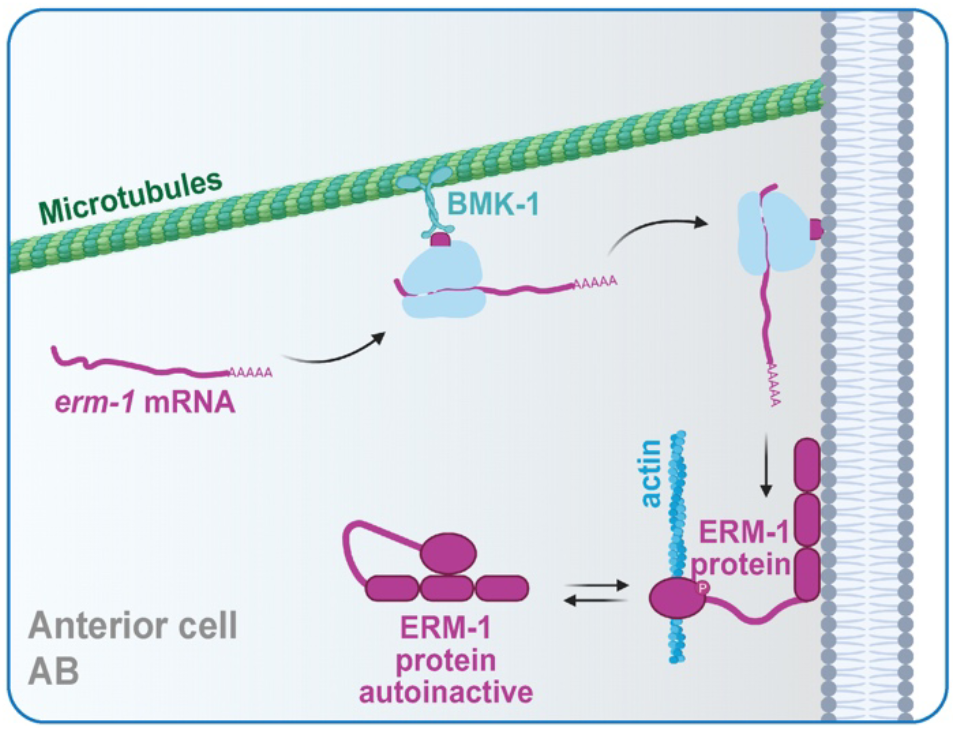
A model of *erm-1*/ERM-1 transport in embryos. **(A)** Model of *erm-1*/ERM-1 translating complex in transit to the membrane via microtubule- and *bmk-1 (klp-14*)-dependent transport. After translation initiation, the exposed N-terminal FERM domain on the nascent peptide directs the entire ribonucleoprotein complex to the membrane in a BMK-1-dependent way. At the membrane, the FERM domain binds PIP_2,_ anchoring it in place.

Many unknowns remain regarding this pathway. We still do not know the identity of the FERM effector protein or if it is even required. The possible links between such an effector and BMK-1 are also unknown. It is unclear whether translation is stalled or slowed in transit or whether multiple rounds of translation occur at the destination. Furthermore, the details of ERM-1’s post-translational migrations are also unclear.

### Why is local translation important for ERM-1 function?

As more membrane proteins are discovered that undergo local translation at membranes, several ideas have emerged to explain why this process occurs. It is possible that local translation stimulates translation in response to external signals and cues, possibly tuning the level of translation appropriately. In the case of very long cells or very rapid processes, local translation may be important to avoid inefficiencies inherent in transporting the formed protein rather than the mRNA. Particularly in cases where mRNAs undergo multiple rounds of translation at the destination, this could result in different rates of protein production (Du et al.,2007; Lécuyer et al., 2009). It is possible that local translation orients nascent proteins so they are perfectly poised to encounter a binding partner or be incorporated into a large complex in the appropriate order relative to other subunits (Engel et al., 2020; Horste et al., 2023; Portz and Shorter, 2018).

All of these scenarios could explain why the ERM-1 proteins rely on local translation for full functionality. However, another idea is that ERM-1 may require local translation to prevent premature auto-inactivation. Although not clearly characterized in *C. elegans*, the human ortholog of ERM-1, Ezrin, exists in either an active, open state or a closed, auto-inhibited state. In the open state, Ezrin’s FERM domain associates with the plasma membrane, its C-terminal ERMAD domain associates with the actin cytoskeleton, and a key T567 residue is phosphorylated. In the closed conformation, N- and C-terminal ends of the protein self-assemble in association with dephosphorylation of T567 and correlated with movement into the cytoplasm (Bretscher et al., 1997; Gary and Bretscher, 1995). Work using phosphomimetic mutants to phenocopy the constitutively active and phosphorylated (T567D) or constitutively inactive, unphosphorylated (T567A) states has shown that Ezrin dynamics can alter the mechanical properties of the cell cortex, the cytoskeletal organization and can impact cancer cell migration and metastasis (Liu et al., 2012; Zhang et al., 2020).

Perhaps local translation facilitates the open, active state by ensuring that ERM family FERM domains form in close proximity to the PIP_2_ phospholipids they bind. Future work will be required to determine how well conserved these processes are in *C. elegans* ERM-1 and if human Ezrin is locally translated.

### Kinesin-5 family members act as transporters of mRNA cargo

It is striking that we identified BMK-1, a Kinesin-5-family member for its impact on mRNA transport. The human ortholog of BMK-1 is KIF11 (Kinesin Family Member 11), also known as Eg6 or Kinesin-5 (Siddiqui, 2002; Yildiz, 2025). KIF11 has been shown to catalyze the transport of β-actin mRNA through interactions with ZBP-1 (Zipcode Binding Protein), a protein that recognizes the “zipcode” sequence within β-actin’s 3’UTR (Biswas et al., 2020; Song et al., 2015). Interestingly, this KIF11-dependent mechanism functions to localize β-actin in fibroblasts and migrating cancer cells. A different kinesin, KIF5A (a Kinesin-1 family member), a myosin (myosin Va also known as MYO5A), and dynein all work to localize β-actin in neurons (Ma et al., 2011; Nalavadi et al., 2012). Previous work has shown that ERM-1 requires the dynein component DHC-1 in L4 and adult intestinal cells (Li et al., 2021). We found that DHC-1 was dispensable for ERM-1 localization in early embryos, and the example of β-actin’s varied and cell-specific transport systems illustrates how such differences can and do arise.

### mRNA mechanisms in early embryos are relevant in other stages

Early embryos harness post-transcriptional mechanisms to drive the earliest stages of cellular differentiation. Studying these mechanisms may yield insight beyond embryos. Indeed, this is the case for *erm-1* as both the maternally inherited and zygotically encoded *erm-1* mRNA transcripts achieve co-translational mRNA localization. Future studies of *erm-1* mRNA and ERM-1 protein together have the potential to link mRNA dynamics and localization to cell morphology and organogenesis.

## MATERIALS AND METHODS

### Maintenance of *C. elegans* strains

*C. elegans* strains were maintained at 20°C in nematode growth medium (NGM: 3 g/l NaCl; 17 g/l agar; 2.5 g/l peptone; 5 mg/l cholesterol; 1 mM CaCl_2_; 1 mM MgSO_4_; 2.7 g/l KH_2_PO_4_; 0.89 g/l K_2_HPO_4_) seeded with OP50 *E. coli* following standard protocols (Brenner, 1974), unless specified otherwise.

### Strains used in this study

| Name | Genotype | Ref |
| --- | --- | --- |
| N2 | Wild type |  |
| LP306 | *cpls53 II; unc-119(ed3) III* | (Heppert et al., 2016) |
| BOX213 | *erm-1(mib15[erm-1::eGFP]) I* | (Ramalho et al., 2020) |

### Embryo permeabilization and drug treatment

To expose embryos to small-molecule drug inhibitors, we first generated synchronized populations of permeable embryos by conducting *perm-1* RNAi knockdown on gravid hermaphrodite mothers. Synchronized L1 hermaphrodites were plated on NGM with IPTG/CARB <concentrations> plates containing *perm-1* RNAi *E*.*coli* concentrated 10x. After ~72 hours, gravid hermaphrodites were collected in M9 and washed twice or until supernatant was clear. Animals were then resuspended in 2 mL of S buffer and transferred to a glass Dounce homogenizer. The hermaphrotide mothers were homogenized, releasing embryos by performing ~5x dounces in a dounce homogenizer and a size “B” pestle. Homogenized hermaphrodites and released embryos were transferred to a microcentrifuge tube and 150 μM Nocodazole in DMSO was added and incubated for 15 minutes at 20°C. Supernatant was removed from embryos, and cooled -20°C methanol was added for fixation. The resulting permeabilized embryos were then used in the standard smFISH protocol.Notably, centrifugation of permeabilized embryos was done at 200 rcf (instead of 2k rcf) x ~1-2 min to prevent the loss of fragile embryos. Plates were stored at 4°C for up to 30 days until used. The 100% embryonic-lethal *pop-1* RNAi was used as a control to test efficacy for each batch of plates.

### RNAi gene knockdown

Synchronized L1 hermaphrodites prepared as described above were plated on NGM/OP50 plates and grown at 20°C for 24-48 hrs, then harvested in M9. Worms were washed twice with M9 until clear. After the final wash, remaining M9 was removed, and animals were seeded on NGM with IPTG/CARB plates containing the *E. coli* strain with the RNAi plasmid of interest at concentrated 10x (see Table S3). Worms were grown at 20°C until gravid (total of 72 hrs). Animals were then collected with M9, washed twice, and prepared for smFISH imaging following the smFISH protocol above.

### smFISH

smFISH experiments were performed as described in (Parker et al., 2021; Spike et al., 2025). Custom smFISH probes for *set-3* and *erm-1* targets were designed using Stellaris RNA FISH Probe Designer (Biosearch Technologies; www.biosearchtech.com/stellaris-designer; version 4.2) and labeled with CalFluor 610 or Quasar 670 (Biosearch Technologies). Hybridization steps used 100 μL of hybridization buffer solution with probes (99 μl Stellaris RNA FISH Hybridization Buffer, 11 μl deionized formamide, 2 μl of 1/20 diluted smFISH probes at 25 μM) and incubated at 37°C for 24-48 hours in a thermoshaker until imaged. After washing and DAPI staining steps, embryos were mounted on a slide with an equal volume of VectaShield antifade (VECTASHIELD® Antifade Mounting Medium; H-1000-10). smFISH image stacks were acquired on a Photometrics Cool Snap HQ2 camera using a DeltaVision Elite inverted microscope (GE Healthcare), with an Olympus PLAN APO 60× (1.42 NA, PLAPON60XOSC2) objective, an Insight SSI 7-Color Solid State Light Engine and SoftWorx software (Applied Precision) using 0.2 μm *z*-stacks. Representative images were deconvolved using Deltavision (SoftWorx) deconvolution software. Representative images were further processed using FIJI and FIJI macros (https://github.com/erinosb/Nishimacros) (Schindelin et al., 2012). All experiments were performed using the symmetrically abundant *set-3* mRNA as a negative internal control for mRNA abundance and subcellular localization. A minimum of 15 embryos were imaged across 2-4 reps or each genetic condition and cell stage.

### DeltaVision image capture, processing and analysis

All images included in this manuscript were captured in a DeltaVision Elite inverted microscope (GE Healthcare), using a Photometrics Cool Snap HQ2 camera and an Olympus PLAN APO 60× (1.42 NA, PLAPON60XOSC2) objective, an Insight SSI 7-Color Solid State Light Engine and SoftWorx software (Applied Precision). Single-color capture was achieved using a beam Splitter suitable for DAPI, FITC, Alexa 594, CY5 and GFP/mCherry and filters with the following single-pass emission settings (DAPI = 435/48, CFP = 475/24, GFP/FITC = 525/48, YFP = 548/22, TRITC = 597/45, mCherry/Alexa594 = 625/45, Cy5 = 679/34). Images were collected using DataVision SoftWorx with identical exposures across any given dataset. Sampling every 0.2 μm in the z-direction. Representative images were deconvolved using DataVision SoftWorx image analysis software and processed using FIJI. At least 10 embryos were imaged for each condition, for each dataset.

### Embryo segmentation, membrane masking and smFISH spot detection

Image analysis was performed prior to deconvolution using an in-house Python pipeline, WormLib, that incorporates cellpose (Pachitariu and Stringer, 2022; Stringer et al., 2021), BigFish (from of fish-quant v2) (Imbert et al., 2022) and other custom-built analysis functions for analysis in *C. elegans*. Embryos were first segmented using cyto2 cellpose model on 2D brightfield images to determine embryo boundaries from the background or other cellular debris in the field of view. Membrane masking was performed using a membrane marker strain (*ph::gfp*) to capture the exact position of membranes. Canny filter with parameters: sigma=0.8, low_threshold=0.08, high_threshold=0.15) were used to mask areas of the membrane using intensity. Images with inaccuracies in embryo segmentation or membrane masks were removed. Spot detection using bigFish was restricted to the embryo regions. mRNA spots were identified in 3D using voxel size of 1448, 450, 450 nm and spot radius of 1409, 340, 340 and 1283, 310, 310 nm for Quasar 670 and Cal Fluor Red 610 channels, respectively. The voxel size and spot radii were optimized for accuracy and consistency.

## ARTIFICIAL INTELLIGENCE (AI) STATEMENT

The authors acknowledge the use of AI to: (1) develop and debug software for analysis of smFISH data; (2) inform and fact check statistical analyses performed using R; (3) complement the use of PubMed for identifying published work potentially informative for the analyses herein; and (4) improve grammar and clarity in the written document.

## ACCESSIBILITY STATEMENT

The colors in microscopy images and plots were selected following guidelines for accessibility. We opted for magenta/green combinations in microscopy images and the viridis and tol packages in R.

## ACKNOWLEDGEMENTS

This work utilized resources from the University of Colorado Boulder Research Computing Group, which is supported by the National Science Foundation (awards ACI-1532235 and ACI-1532236), the University of Colorado Boulder, and Colorado State University. This work utilized microscopy resources from NIH grant 1S10 OD025127 and support from the CSU Microscope Imaging Network. Some strains were provided by the Caenorhabditis Genetics Center, which is funded by National Institutes of Health Office of Research Infrastructure Programs (P40 OD010440). Some figure elements were created in BioRender. We are grateful to Dr. Carol Wilusz, Dr. Tom LaRocca, Dr. Tim Stasevich, Dr. Matthew Taliaferro, Dr. Mike Boxem, Dr. Suzan Ruijtenberg, and Mette Schroeder for valuable discussion and feedback on this project.

## COMPETING INTERESTS

The authors have no competing or financial interests.

## AUTHOR CONTRIBUTIONS

Conceptualization: N.T., E.O.N.; Methodology, formal analysis, and investigation: N.T., K.C., E.O.N.; Writing, review, and editing: N.T., K.C., E.O.N.; Supervision, project administration and funding acquisition: N.T., E.O.N.

## FUNDING

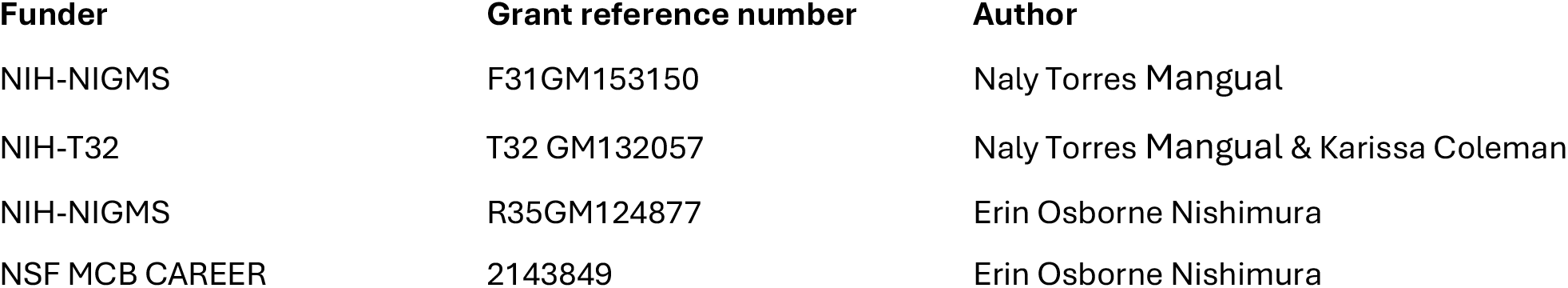

## Notes

**Funding:** This work was supported by the National Institute of General Medical Sciences (F31GM153150, R35GM124877, T32GM132057) and the National Science Foundation (2143849)

### Competing Interest Statement

The authors have declared no competing interest.

